# Distinct neural oscillations predict covert switching between stable and unstable cognitive representations

**DOI:** 10.1101/2025.05.19.654819

**Authors:** Judith Sattelberger, Hamed Haque, Liu Mengxing, Jaana Simola, J Matias Palva, Michael M. Halassa, Satu Palva

**Affiliations:** Neuroscience Center, Helsinki Institute of Life Science, University of Helsinki, FI-00014 Helsinki, Finland; BioMag Laboratory, HUS Medical Imaging Centre, Helsinki University Central Hospital, FI-00029 HUS, Finland; School of Psychology and Neuroscience, University of Glasgow, G12 8QB Glasgow, UK; Department of Neuroscience, Tufts University School of Medicine, Boston, MA 02111, USA; Faculty of Educational Sciences, University of Helsinki, FI-00140 Helsinki, Finland; Department of Neuroscience and Bioengineering (NBE), Aalto University, FI-02150 Espoo, Finland

**Author notes:** Correspondence should be addressed to Judith Sattelberger or Satu Palva.

**Keywords:** cognitive control, oscillations, task switching, synchrony, WCST

## Abstract

Cognitive flexibility is the ability to keep or update contextual cognitive representations. The underlying neural mechanisms and how it is influenced by individual sensitivity to feedback are not known. We measured cognitive flexibility under uncertainty with the Wisconsin Card Sorting Task and brain activity with magnetoencephalography (MEG). Using a behavioral sequential learning model, we show that variability in cognitive flexibility is predicted by individual reward and punishment sensitivity to feedback and anticipation and exploration tendencies. Uncovering their brain oscillatory signatures showed that individual learning speed is predicted by suppressed alpha-beta (7–32 Hz) and increased broad-band gamma (52–96 Hz) amplitudes along with concurrent large-scale alpha (7–13 Hz) desynchronization. Importantly, these oscillatory sub-processes were explained by individual levels of distinct feedback sensitivities. These results provide a novel account on neural sub-computations underlying flexible contextual switching between stable and unstable cognitive representations that predict the individual speed of rule learning.

## Introduction

Cognitive flexibility is a key component of cognitive control alongside top-down attention and inhibition, enabling us to quickly adapt behavior to an unpredictable and changing environment. This executive function varies widely between individuals and over the lifetime (Rhodes, 2004). At its core, cognitive flexibility allows us to make decisions under uncertainty, based on cognitive rule representations whose stability depends on feedback from the environment. For example, identifying the right key of a set of unmarked keys to open a door requires remembering (i.e., using a mental map of) which key opens which door, but because such mental maps may be uncertain, a trial-and-error approach can augment the search until the right key is identified by the opening of the door. The internal representation of which key has to be used remains unstable until the door finally opens. A fundamental question is how such cognitive flexibility for switching between cognitive rules (or keys), leading to stable and unstable cognitive representations, is achieved in the brain. The ability to adapt to changing task rules, even when they are not explicitly instructed, is called task switching and has often been studied with the Wisconsin Card Sorting Test (WCST) in humans (Barcelo, 2003; Berg, 1948), monkeys (Kuwabara et al., 2014; Mansouri et al., 2006) and recurrent neural networks in computational (Liu & Wang, 2024). Performance in the WCST depends on multiple components, namely global and local switching, error monitoring and reward processing (Gamboz et al., 2009; Miyake et al., 2000; Nyhus & Barceló, 2009), but only one study has tried to distil the impact of those sub-processes on interindividual performance variability in the WCST using a computational model (Bishara et al., 2010).

In the brain, switching between cognitive representations depends critically on activation of the prefrontal cortex (PFC) (Buckley et al., 2009), in line with the PFC’s general role in cognitive control tasks (Bissonette et al., 2013; Panikratova et al., 2020; Robbins et al., 1996), but also on activity at posterior parietal and occipital cortices (Periánez et al., 2004; Wang et al., 2001). Research in non-human animals has further shown that the mediodorsal thalamus and its interactions with the PFC are key to enabling cognitive flexibility (Lam et al., 2025; Rikhye et al., 2018), especially under task uncertainty (Zhang et al., 2025). Accumulating evidence indicates that brain rhythms – neuronal oscillations – are fundamental for temporal coordination of neuronal processing in a variety of cognitive functions (Siegel et al., 2012; Thut et al., 2012; Vinck et al., 2023; Voytek & Knight, 2015).

Two decades of work leveraging electroencephalography (EEG) to study overt task switching in relation to sensory representations have revealed theta (θ, 3–8 Hz) band oscillations in the PFC or the midline θ activity to play a pivotal role in cognitive control (Duprez et al., 2020; Muralidharan et al., 2023) including conflict monitoring (Riddle et al., 2020; Sauseng & Liesefeld, 2020) and anticipation (Cooper et al., 2015; Khan et al., 2025). There is also some evidence for the role of gamma (γ, 30–100 Hz) (Proskovec et al., 2019), parietal alpha (α, 8–12 Hz) (Foxe et al., 2014; Verstraeten & Cluydts, 2002), and beta band (β, 13– 30 Hz) (Cunillera et al., 2012; Gladwin et al., 2006) amplitudes in task-switching performance although this evidence is less consistent. Recent evidence, importantly, suggests that low-frequency brain oscillations could facilitate switching between the mental states associated with behavioral stability and flexibility in cognition (Ericson et al., 2025). In parallel, switching between conscious sensory representations has been associated with anterior–posterior network communication and anterior information differentiation (Canales-Johnson et al., 2023).

We hypothesized that adaptations in oscillation dynamics and their large-scale networks could be a neural mechanism supporting task switching in the WCST. So far, no studies have resolved how these neural modulations might enable the switching between stable and unstable cognitive representations nor whether these processes would underlie the inter-individual variability in cognitive flexibility performance. Here, we took advantage of the rich local field potential data from magnetoencephalography (MEG) (Baillet, 2017) to study the neural adaptations taking place during the WCST with excellent temporal resolution. Crucially, by applying an individualized source reconstruction approach for every test subject, MEG enables precise anatomical mapping of neuronal sources and their effective inter-areal connectivity. Such synchronization of neural oscillatory signals within as well across oscillatory frequencies has been found to support top-down attentional selection (D’Andrea et al., 2019; Lobier et al., 2018; Rouhinen et al., 2020) and explain individual performance variability in visual working memory (VWM) performance (Mamashli et al., 2021; Palva et al., 2010; Sattelberger et al., 2024; Siebenhühner et al., 2016). Despite the likelihood that dynamic oscillatory network fluctuations would be fundamental for enabling internally guided switching of cognitive rules or tasks based on feedback processing, this has remained uncharted.

In this work, using MEG data, we investigated neural computations underlying the internal switching performance to study the possibility that neural low-frequency oscillations and their large-scale network interactions could provide such a mechanism. To explain the individual variability in cognitive flexibility we implemented a new behavioral sequential learning model with which we estimated the impact of feedback processing on rule learning speed. We then identified which neural signatures of cognitive flexibility and learning speed were explained by sensitivity to feedback and individual anticipation and exploration tendencies.

## Results

In this study, 26 healthy participants completed a computer-based version of the Wisconsin Card Sorting Test (WCST) in which the active sorting rule (out of three possible rules) changed after every fifth trial (Fig. 1A). This task thus allows the separation of rule representations into stable (Stay trials, in which the internal rule remains the same) and unstable (Shift trials, where the internal rule changes) cognitive representations, whereby the subject’s performance across the trial set measures cognitive flexibility. As the subjects were not informed about the rule change interval, the active rule needed to be sorted out from negative (Shift) or positive feedback (Stay), which we analysed with a sequential learning model (Fig. 1B). With this model, we estimated the currently active cognitive rule representations and their stability (rule certainty) based on trial-to-trial task decisions. By combining the subject’s task decisions and feedback across trial sets, we obtained the subject parameters of positive and negative feedback processing. MEG data during rule selection were source-reconstructed using individual anatomical MRIs and collapsed into a cortical parcellation of the Schaefer atlas and filtered with Morlet wavelets between 3-143 Hz. A data-driven approach was then used to compute local and inter-areal oscillation dynamics for all frequencies and for all parcels or parcel-pairs, respectively (Fig. 1C).

**Figure 1.**
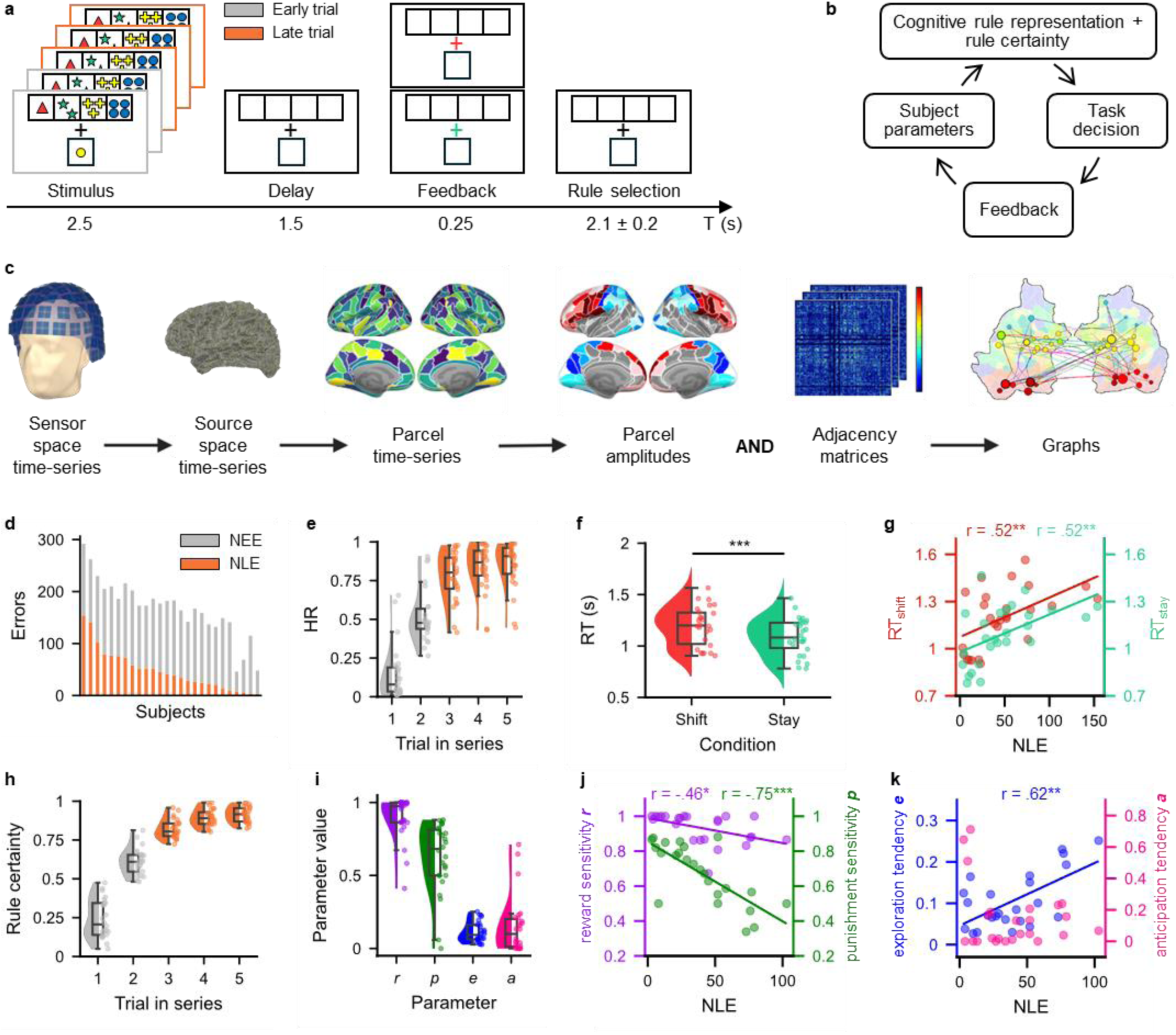
Experimental paradigm and behavioral results. **a.** Schematic of the Wisconsin Card Sorting test. In each trial, a new sample card (bottom row) and the four target cards (top row), were shown, followed by a delay during which the participants were to match the sample card to one of the target cards based on a changing rule. A green (correct) or red (incorrect) feedback cross was shown after the response, followed by a delay that allowed for selection of a cognitive rule for the next trial. **b**. Schematic of the behavioral modelling. In the beginning of each trial, the participant had a cognitive rule representation, based on which they made a task decision to match the card to one of the three categories, leading to positive or negative feedback. Depending on the subject’s individual tendency to respond to feedback as estimated by the subject parameters, this feedback modified the rule representation and increased the certainty of that rule. **c.** Schematic of the MEG analysis. Cortical sources were reconstructed from broad-band MEG time-series data, collapsed into cortical parcels and used for estimating oscillation amplitudes and phase-synchronization between parcels. **d.** Number of early and late errors (NEE and NLE) by subjects (N = 26). **e.** Averaged HRs for every trial in a rule set. The half violin plots and boxplots show the distribution and median values of HRs, and the dots the subjects’ means. **f.** The difference in RTs for shift and stay conditions between subjects. Asterisks indicate significant differences (Wilcoxon signed-rank test, *p < 0.05, **p < 0.01, ***p < 0.001). **g.** Scatterplot of the correlations of RTs with NLE across participants. Colored fitting lines indicate the significant correlation of NLE with RT_shift_ (red) and RT_stay_ (green). Asterisks indicate a significant correlation (Pearson’s r). **h.** Mean rule certainty by trial order and across subjects. **i.** Parameter values across subjects. **j-k.** Correlations of all subject parameters with NLE across participants (N = 24). Dots represent each subject’s value in reward sensitivity (purple), punishment sensitivity (green), exploration tendency (blue) and anticipation tendency (pink), and colored fitting lines indicate significant correlations of the respective subject parameter with NLE.

### Behavioral performance

In the WSCT task, the participants needed to match cards according to three rule-categories: shape (circle, star, cross, or triangle), color (blue, yellow, red, or green), or number of items (one to four). The category by which to match changed on every fifth trial unknown to the participants, so that the rule needed to be deduced based on feedback on whether the preceding selected rule was correct (green cross) or incorrect (red cross). Consequently, if the selected rule was correct, the participant could “stay” at the current rule during the next trial, while if it was incorrect they were to “shift” to a different rule. This allowed us to measure the neural signatures of stable (Stay trials) vs. flexible / changing (Shift trials) cognitive rule representations across participants and further, to investigate how individual differences in the ability to switch between representations depended on the participants’ sensitivity to feedback. To characterize the accuracy and rapidness of switching between stable and new flexible representations and its variability across participants we categorized incorrect responses during the first two trials of a rule series as early errors (EE) and all subsequent errors as late errors (LE) based on the required number of trials needed to deduce the rule from feedback alone. Calculating the number of LE (NLE) allowed us thus to assess the participant’s actual learning speed beyond chance rate. Across participants, there was a high variability in NLE establishing the individual differences in cognitive flexibility, i.e., the tendency to maintain stable or switch to new cognitive representations under uncertainty (Fig. 1D).

However, generally, participants’ higher number of EE (NEE; M = 119.3, SD = 27.8) than NLE (M = 47.9, SD = 38.9), reflected their rule learning across a rule set of five trials. This was further visible in HRs increasing with every trial within a rule set, showing a clear gradient in HRs from early to late trials (Fig. 1E). RTs were on average 102 ms longer in shift than stay trials (Fig. 1F; *T*(25) = 6.01, *p* = 2.81e-6; BF10 = 6879.316), meaning that participants reacted more slowly after having received negative feedback. Additionally, both RT_shift_ (r = 0.52, p = 0.0065, BF10 = 8.110) and RT_stay_ (r = 0.52, p = 0.0063, BF10 = 8.398), but not their difference RT_shift_ - RT_stay_ (r = 0.05, p = 0.7963, BF10 = 0.251), were positively correlated with NLE across participants (Fig. 1G). Differences in behavior were neither confounded by age [HR (r = - 0.13, p = 0.514, BF10 = 0.298), NLE (r = 0.05, p = 0.827, BF10 = 0.249), RT_shift_ (r = 0.11, p = 0.573, BF10 = 0.283), RT_stay_ (r = 0.16, p = 0.422, BF10 = 0.331)] nor sex [HR (T(24) = -1.281, p = 0.212, BF10 = 0.664), NLE (T(24) = 1.220, p = 0.234, BF10 = 0.629), RT_shift_ (T(24) = 1.159, p = 0.258, BF10 = 0.597), RT_stay_ (T(24) = 1.382, p = 0.180, BF10 = 0.730)].

With our behavioral model, we first estimated the trial-to-trial rule certainty by creating an attention-weight-vector that represents the participant’s chosen rule in the previous trial (Bishara et al., 2010), whereby the rule certainty is calculated as the inverted Jensen-Shannon distance between weight vectors. This allowed us to investigate trial-to-trial changes associated with the participant’s certainty (stability) of the rule representation across a series of trials within a given rule set (Fig. 1H), which overall increased with every trial of a rule series from a mean value of 0.242 at the first trial to 0.913 at the fifth trial (F(4) = 297.209, p < .001, BF10 = 8.226e+55). To then investigate how cognitive flexibility was driven by differences in the sensitivity to positive – reward – or negative – punishment – feedback at the single-trial level for each individual, we implemented a sequential learning model with which we estimated the subject-wise parameters for reward sensitivity ***r*** and punishment sensitivity ***p*** measuring error monitoring as in Bishara et al. (2010). In addition, as new behavioral measures, we estimated the exploration tendency ***e*** and anticipation tendency ***a*** which assessed the effect of rule shifting despite preceding positive feedback and the subjects’ ability to predict an upcoming rule change after 5 trials, respectively (Fig. 1I). We then used multiple linear regressions to reveal the explanatory value of these parameters on HRs, RTs and NLE. The model was significant for HRs (F(4,21) = 62.703, p < .001, R² = 0.923, BFM = 3.852) and NLE (F(4,21) = 70.818, p < .001, R² = 0.931, BFM = 29.342), but not RT_stay_ (F(4,21) = 1.619, p = 0.207, R² = 0.236, BFM = 0.500) or RT_shift_ (F(4,21) = 1.593, p = 0.213, R² = 0.233, BFM = 0.499), establishing that the accuracy but not speed depends on individual feedback sensitivity. The regression equation for HR was thus HR = 0.277 + (0.141***r***) + (0.315***p***) - (0.361***e***) + (0.430***a***) where all parameters but ***r*** were significant (***r***: p = 0.108, BFi = 1.212; ***e***: p = 0.017, BFi = 9358.513; ***a:*** p < 0.001, BFi = 3.205; ***p***: p < 0.001, BFi = 2.909e+06). The number of late errors were explained by NLE = 153.2 – (62.5***r***) – (97.5***p***) + (191.8***e***) – (62.9***a***) with all parameters significant (***r***: p = 0.018, BFi = 7.478; ***p***: p < 0.001, BFi = 9733.338; ***e***: p < 0.001, BFi = 231.102; ***a***: p < 0.001, BFi = 589.644).

Responsivity to positive and negative feedback positively covaried within subjects (Pearson‘s r = 0.41, p = 0.047, BF10 = 1.634), so that participants with a higher reward sensitivity tended also to be more sensitive to punishment. While higher sensitivity to either form of feedback was associated with higher learning speed, however, punishment sensitivity was a much stronger predictor of NLE across subjects (Fig. 1J; ***r***: Pearson‘s r = -0.46, p = 0.023, BF10 = 2.865; ***p***: Pearson‘s r = -0.75, p < 0.001, BF10 = 1028.328), demonstrating the distinct effect of positive and negative feedback on improvements in the behavioral outcome. Additionally, subjects with a larger exploration tendency had a higher NLE (Fig. 1K; ***e***: Pearson‘s r = 0.62, p = 0.001, BF10 = 35.356), whereas the tendency to anticipate upcoming rule changes did not significantly impact learning speed (***a***: Pearson‘s r = -0.28, p = 0.185, BFi = 0.581).

### Oscillation amplitudes predict individual differences in feedback processing

To investigate the neural mechanism underlying these behavioral effects, we computed oscillation amplitudes from individually source-reconstructed MEG data (Fig. 1C). We first computed these for the Shift and Stay trials focusing analysis on the rule selection period, during which participants decided whether to maintain the current cognitive representation for the rule or update it for a new rule. We compared Shift and Stay trials to the pre-stimulus baseline and then calculated significant differences between the two conditions (Fig. 2A, Wilcoxon signed-rank test, p < 0.05, q = 0.08). Both Shift and Stay conditions were characterized by a strong sustained low-theta band (θ, 3–6 Hz) oscillations and a concurrent suppression of alpha (α, 7–13 Hz) and beta (β, 13–32 Hz) band oscillation amplitudes throughout the rule selection period. In addition, the Shift condition was also associated with an increase in broad-band gamma band (γ, 52–96 Hz) band amplitudes. These amplitude modulations were significantly stronger in Shift than Stay trials and had medium to strong effect sizes (Supp. Fig. 1A, θ: p = 2.03e-34, r = 0.87; α: p = 6.88e-16, r = 0.57; β: p = 3.28e-26, r = 0.75; γ: p = 9.65e-31, r = 0.82), indicating that these modulations were associated with switching of cognitive representations. Differences in θ-band oscillation amplitudes were widespread across frontal, temporal and visual cortices, whereas α and β amplitudes were suppressed over parietal areas with a concurrent amplitude increase in the left lateral (sub-)orbital sulcus and the bilateral primary visual areas, largely overlapping between the two frequency bands, leading us to combine amplitude effects in these frequency bands (Supp. Fig. 1B). Increases in γ amplitudes were located in the left cingulate cortices, precuneus, insula and orbitofrontal cortex and the parcels of V1 (right calcarine sulcus and cuneus).

**Figure 2.**
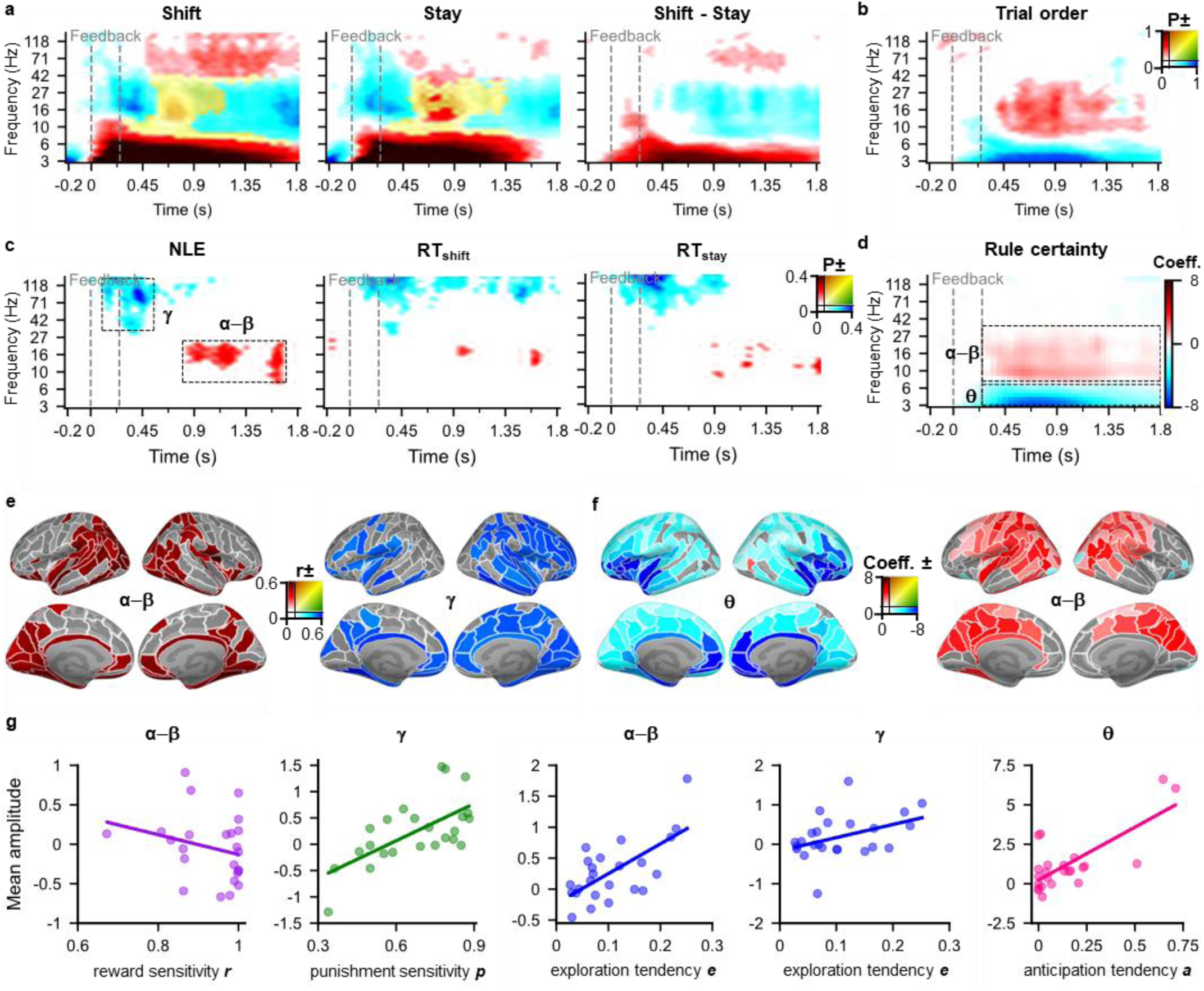
Oscillation amplitude effects. **a.** Group-averaged oscillations amplitudes for the Shift and Stay trials compared to the baseline, and their contrast (N = 26). Color indicates the fraction of parcels (brain regions) (*P*) with a statistically significant effect (Wilcoxon signed-rank test, p < 0.05, FDR corrected, q = 0.08). The dashed black box indicates the time-frequency (TF) bins of interest during rule selection. **b.** The fraction of parcels whose amplitude significantly increased or decreased across a rule set (Spearman’s r, p < 0.05, FDR corrected, q = 0.08). **c.** Correlation of parcel amplitudes with NLE, and the Shift and Stay RTs (Spearman’s r, p < 0.05, FDR corrected, q = 0.08). **d.** Oscillation amplitudes explained by increasing rule certainty within a trial series, where colors indicate the coefficients of parcels with a significant model fit (Linear Regression, p < 0.05, FDR corrected, q = 0.08). **e.** Brain parcels with a significant correlation with NLE. Color indicates the correlation coefficients. **f.** Brain parcels with a significant increase in rule certainty across trials. Color indicates the regression coefficients. **g.** Relationship of mean amplitudes of Shift and Stay trials over all parcels with a significant model fit and the subject parameters. One circle represents one subject and lines indicate the fit across subjects (N = 24).

Next, to study whether oscillation amplitude modulations would change as participants learned the currently active rule, we computed the correlation of oscillation amplitudes with trial order, i.e., the position of the trial within a rule set (Fig. 2B, Spearman’s r, p < 0.05, q = 0.08). Participants showed smaller θ amplitudes between 0.9–1.8 s after feedback, and increased α−β amplitudes (0.5–1.35 s after feedback) as the rule set progressed, indicating that the θ−β amplitudes underwent a spectral transition from rule shifting towards staying at the correct rule and indicating that these modulations were associated with increased stability of cognitive representations.

To further understand the neural basis of individual variability in behavioral measures of cognitive flexibility, we investigated how the condition-averaged amplitudes would predict differences across individuals in behavior and how they would be explained by the subject-specific parameters from the sequential learning model. Individual variability in switching behavior, especially variability in NLE, but also in RT_shift_ and RT_stay_, correlated positively only with α−β and γ amplitudes while there was no effect in the θ amplitude (Fig. 2C). We next used the trial-to-trial rule certainty as a regressor for oscillation amplitudes to understand the effect of rule representation stability on local neural activity. Higher rule certainty was associated with smaller θ band and larger α−β band amplitudes (Fig. 2D) similarly to the trial order correlation indicating that the spectral amplitude transition across a trial series was related to increased rule certainty. Positive correlations with NLE were found in the striate cortices, posterior parietal cortex (PPC), and in the medial PFC including insular and suborbital cortices (Fig. 2E). In addition, NLE was correlated negatively with broad-band γ amplitudes arising from the parcels in the visual system, the intraparietal sulcus (IPS) of the PPC, and from both the lateral and medial PFC including anterior cingulate cortices. Thus, participants with fewer NLE had smaller α−β amplitudes over the PPC and larger γ-band activity. The spectral changes associated with increasing rule certainty were most prominently seen in the PFC in the θ band and over the PPC in the α−β band (Fig. 2F).

To then investigate whether subject-wise differences in feedback sensitivity could explain different sub-processes underlying the neural signatures of behavior-oscillation amplitude correlations, we first correlated the amplitudes with all frequencies (Supp. Fig. 2B) and then pinned down the effects in the frequencies we just established as relevant with subject-specific model parameters (Fig. 2G). The most robust effect of α−β band related to rule updating and rule certainty was best explained by the reward sensitivity ***r*** so that subjects that were more sensitive to positive feedback showed more decreased α−β amplitudes. In contrast, we found distinct neural signatures to be explained by sensitivity to other feedback. The punishment sensitivity ***p*** and exploration tendency ***e*** were predicted robustly by a sustained γ-band (43– 143 Hz, 0.5–1.8 s) amplitude increase, and ***e*** additionally by a brief transient increase in α−β amplitudes. In contrast, anticipation tendency, ***a***, was predicted by a sustained θ (3–8 Hz, 1–1.8 s) amplitude increase. These feedback-specific neural processes also displayed distinct anatomical distributions across the cortex, however mirroring the effect localizations of Shift-Stay amplitude differences in the respective frequency bands (Supp. Fig. 2C). Hence, we found specific neural oscillatory subprocesses to underlie covert task switching under rule uncertainty and to reflect differences in feedback processing.

### α desynchronization predicts individual behavioral cognitive flexibility

To resolve whether oscillation-based functional connectivity could reflect cross-cortical routing of information processing to enable cognitive flexibility, we used a measure of inter-areal phase synchronization that indicates the statistical temporal regularities between brain areas. To this end, we computed inter-areal phase synchronization with the imaginary phase-locking value (iPLV) between all cortical parcels and for all frequencies, separately for Shift and Stay trials against the baseline and then the difference between the two conditions (Fig. 3A, Wilcoxon signed-rank test, p < 0.05, q = 0.0067). Both the Shift and Stay conditions were characterized by strong θ-band synchronization that was sustained throughout the rule selection period and by a later onset α-band desynchronization, both of which were stronger for the Shift than Stay condition. However, trial order was only correlated with a decrease in θ synchronization (Fig. 3B), indicating that the synchronization decreased as the rule set progressed from Shift towards Stay trials and i.e. that synchronization was specifically related to switching of the internal rule representation. The largest differences between Shift and Stay conditions had a medium effect size in the θ band (Supp. Fig. 3, Wilcoxon signed-rank test; p = 0, r = 0.44) and a small effect size in the α band (p = 3.34e-154, r = 0.18). As for the local oscillation amplitudes, we investigated whether mean strength of phase synchronization correlated across individuals with NLE, RT_shift_ and RT_stay_. A smaller NLE, i.e., increased learning speed, was predicted by α-band desynchronization, however, the network synchronization did not predict RTs for Shift and only weakly for the Stay condition (Fig. 3C).

**Figure 3.**
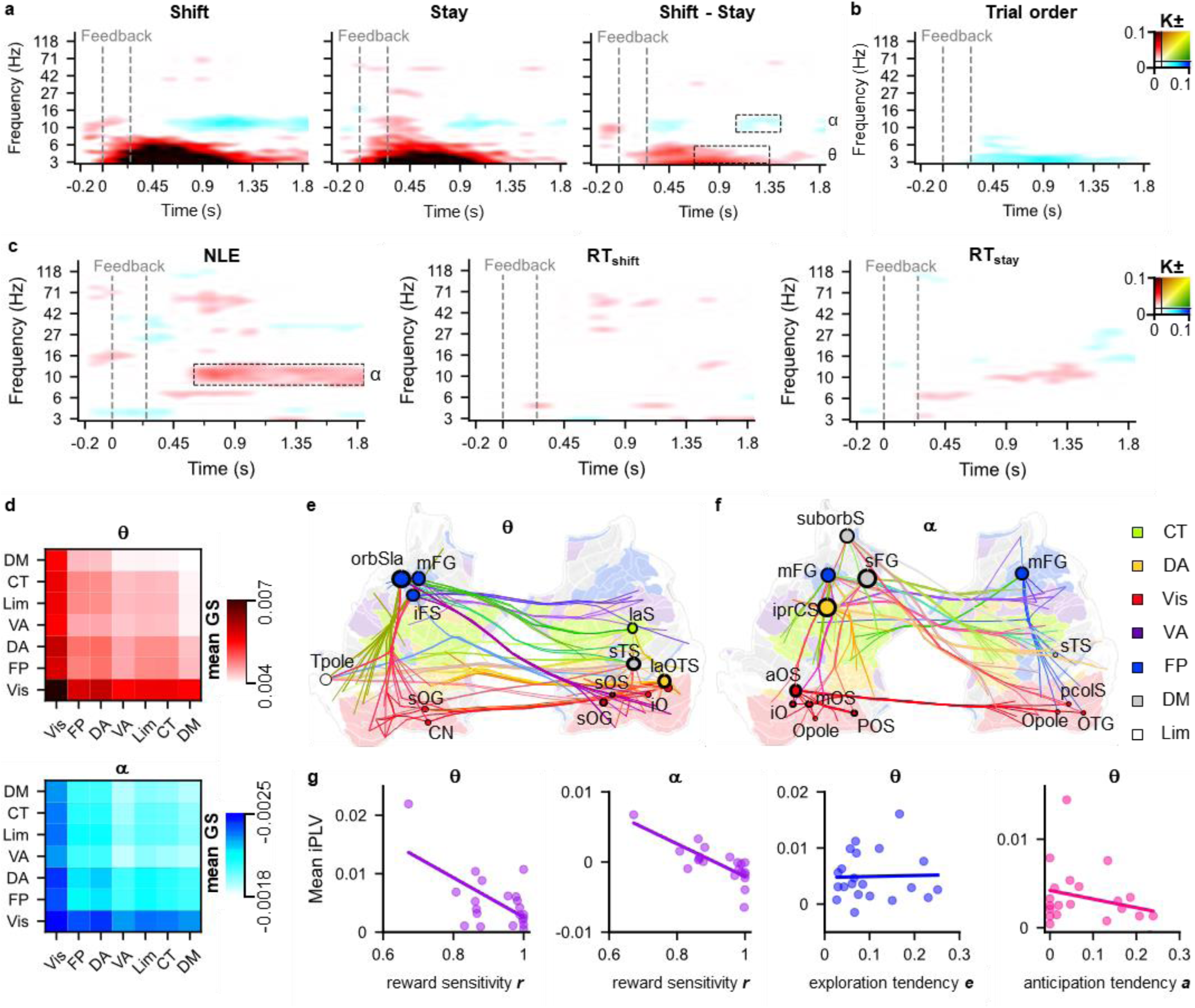
Phase synchrony (iPLV) results for Shift and Stay conditions, the correlations with performance and the model fit. **a.** Interareal phase-synchronization for the Shift and Stay trials compared to the baseline and their contrast Shift-Stay (Wilcoxon signed-rank test, p < 0.05, FDR corrected, q = 0.0067) across participants (N = 23). Color indicates the connection density (*K*) of significant connections. **b.** Trial series effect showing the fraction of edges whose strength significantly increased or decreased across a rule set (Spearman’s r, p < 0.05, FDR corrected, q = 0.0067). **c.** Correlation of the strength of phase synchronization for each edge with NLE, and RTs by frequency band (Spearman’s r, p < 0.05, FDR corrected, q = 0.0067). **d.** Mean graph strength (mean GS) averaged over the functional networks in the θ band (top) and α band (bottom). **e.** θ-band (3-6 Hz, 0.7-1.3 s) matching graph showing the vertices with the highest degree and the 150 edges with the strongest phase-synchronization. Graphs are visualized on a flattened cortex where the color refers to the functional Yeo-network. Lines indicate the 200 edges with highest degree and are colored according to the connected parcels and networks. Spheres and annotations indicate nodes, with radii proportional to their degree, and borders proportional to their eigenvector centrality. **f.** α-band (7-13 Hz, 0.6-1.8 s) matching graph showing the network of the nodes with the largest degrees and strongest edges correlating with NLE. **g.** Model fit for each subject parameter with the mean edge strength of Shift and Stay trials (N = 21). The color shade expresses the strength of the regression coefficients (Coeff., linear regression, p < 0.05, FDR corrected, q = 0.0068).

The differences of Shift-Stay conditions in the θ- and α-bands were driven by functional connections within the visual system and its connections to other networks (Fig. 3D). Visualization of the θ network architecture with the 200 strongest edges and nodes with the largest degrees (i.e., the number of edges with other nodes) revealed major hubs in the PFC (including the middle frontal gyrus (mFG) and inferior frontal sulcus (iFS)) coupled with nodes of the visual system, such as the superior occipital gyrus (sOG) and cuneus (CN) and lateral occipitotemporal sulcus (laOTS) (Fig. 3E). The visuo-frontal network architecture was thus well supporting the idea of synchronization enabling top-down visual selection of correct rule representation. Similarly, the α-band network related to learning speed comprised mostly frontoparietal edges between the suborbital sulcus (suborbS), superior and medial frontal gyri (sFG and mFG), inferior precentral sulcus (iprCS), sTS, anterior occipital sulcus (aOS), mid-occipital sulci (mOS), iO, the bilateral occipital poles (Opole), parieto-occipital sulcus (POS), pcolS, and the occipitotemporal gyrus (OTG) (Fig. 3F).

To understand how these network effects related to individual differences in feedback processing, we then carried out analysis at the single-trial level with the subject-specific model parameters which showed that α desynchronization was predicted by increased reward sensitivity ***r*** but not by other parameters i.e. punishment sensitivity ***p***, anticipation tendency ***a*** or exploration tendency ***e,*** while in contrast θ-band synchronization (Fig. 3G) characteristic to the Shift-Stay difference was explained by anticipation and exploration tendencies together with reward sensitivity.

### Nested coupling of θ phase to α, β, and γ amplitudes predicts task-switching performance

In the previous analysis, we had established that local α to γ amplitudes and concurrent θ synchronization are relevant to endogenous task-switching. To study whether integration of these distinct processes could be achieved via cross-frequency interactions, we computed phase amplitude coupling (PAC) between low frequency concurrent θ oscillation phase and high frequency oscillation amplitudes. Both Shift and Stay conditions were characterized by strong PAC between low to high-θ (3–8 Hz) phase and α−γ amplitudes (9–143 Hz, Fig. 4A) such that θ−γ PAC was stronger in Shift than Stay trials and decreased over the course of a rule series (Fig. 4A–B). Post-hoc correlative analysis revealed that individuals with larger PAC of high-θ to all HFs predicted fewer NLE, RT_shift_ and RT_stay_ (Fig. 4C). θ−γ PAC was found to connect the θ phase of prefrontal parcels with the γ amplitudes of visual parcels (Supp. Fig. 4A), and the same networks predicted fewer NLE and shorter RTs across participants (Supp. Fig. 4B).

**Figure 4.**
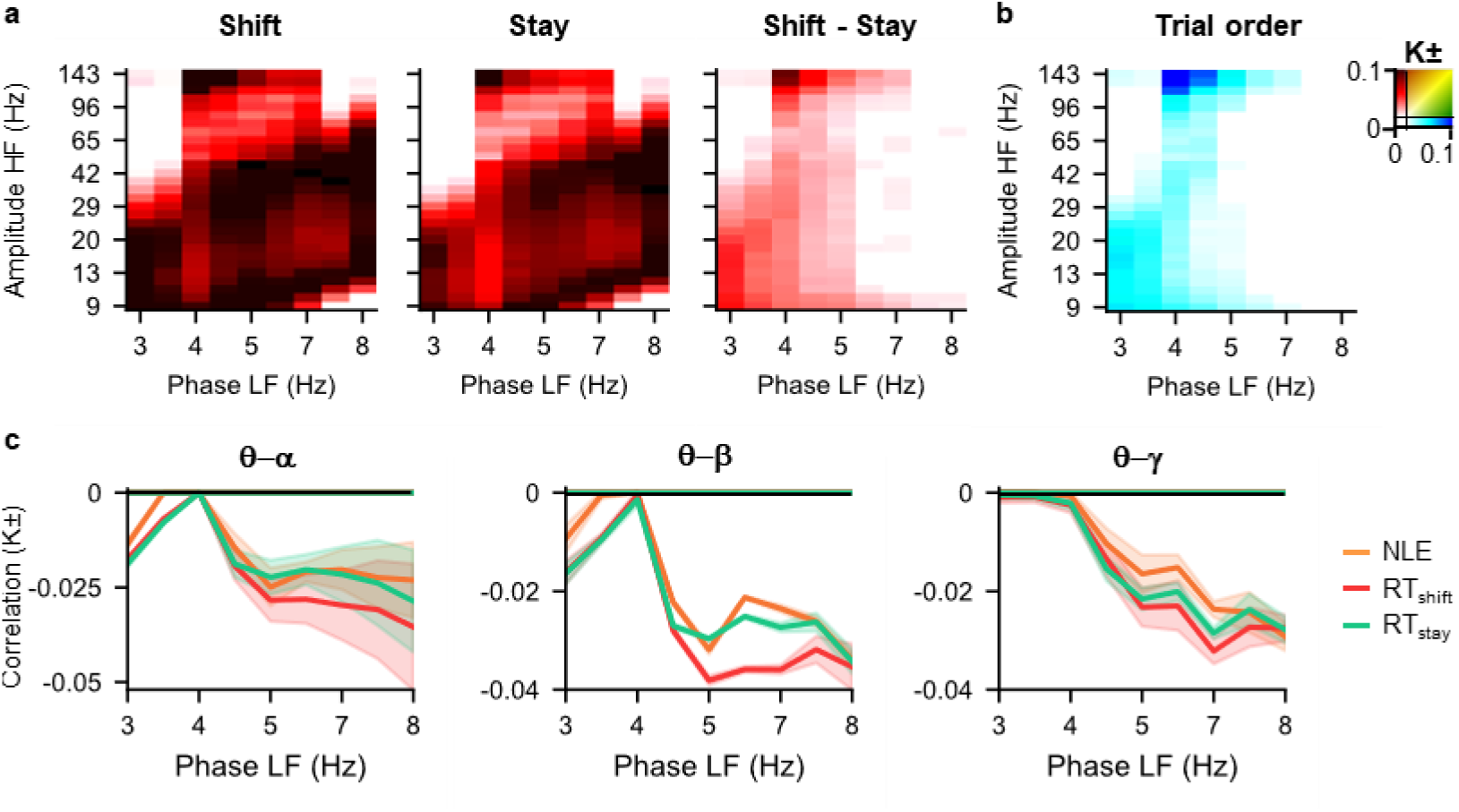
Phase amplitude coupling results for Shift and Stay conditions, trial order, and the correlations with performance. **a.** Phase amplitude coupling (PAC) between low-θ frequencies (LF) and high-α to γ frequencies (HF) during Shift and Stay trials compared to baseline (surrogate statistics, p < 0.01, corrected for spurious connections, q = 0.035) and their contrast (Wilcoxon signed-rank test, p < 0.05, FDR corrected, q = 0.00389) across participants (N = 23). Color indicates the connection density (K) of significant connections. **b.** Trial series effect showing the fraction of edges with significant in- or decrease in PAC across a rule set (Spearman’s r, p < 0.05, FDR corrected, q = 0.00389). **c.** Positive (K+) and negative (K-) tails of the correlations of PAC across frequency-band-averaged HFs with NLE, RT_shift_ and RT_stay_ (Spearman’s r, p < 0.05, q = 0.00281).

## Discussion

Cognitive flexibility is the ability to quickly adapt behavior to changing environmental demands that may require either to keep (Stay with) the current cognitive representation or update (Shift) the cognitive representation to a new one. In the real world, this adaptation often depends on the interpretation of ambiguous instructions and environmental cues with high uncertainty. Individuals vary widely in their ability of such cognitive switching between stable and flexible mental representations as well as on their ability to interpret environmental cue information with high uncertainty as also evidenced in our study by the large heterogeneity across participants in the number of late errors to update to a new rule. The precise neural reasons of these behaviors have remained unknown along with a limited understanding of how feedback processing could influence switching between the internal representations of the contextual rules. Using advanced analysis of MEG oscillation dynamics data and a new computational modelling approach of behavioral performance, we demonstrate that in contrast to previous findings, rather than θ-band oscillations, both local α−β as well as broad-band γ band oscillations and their large-scale networks predict individual variability in the flexible switching between stable and new cognitive representations, each reflecting distinct subprocesses related to feedback processing.

Using a new computational model of subject-wise trial-to-trial rule learning, we show that the subprocesses underlying interindividual variability in cognitive flexibility are explained by differences in positive (reward) and negative (punishment) feedback sensitivity, as well as by exploration tendency, i.e., the tendency to update the cognitive rule representation despite positive feedback. Importantly, this variability was not driven by demographic factors such as sex or age in our sample likely due to the young age of our participants (Rhodes, 2004), verifying that interindividual differences in oscillatory modulations can be expected to reflect actual differences in cognitive flexibility.

For the first time, we further show that oscillation dynamics is a neural mechanism that explains the inter-individual variability in cognitive flexibility. Enhanced broad-band γ-band oscillation amplitudes and concurrent suppression of α−β (13–32 Hz) band amplitudes and synchronization predicted the ability to switch between cognitive representations and the rapidity thereof. The computational modelling of individual performance at the single-trial level established that α- and β-band amplitude suppression predicted more rapid cognitive flexibility as a result of subjects’ increased reward sensitivity, while in contrast the behavioral advantage of increased γ amplitudes arose from higher punishment sensitivity. Thus, α−β band amplitude suppression and the increase in γ amplitudes assigned to different neural subprocesses that predicted the variance in cognitive flexibility across individuals.

The importance of α−β-band amplitudes in switching between stable and new cognitive representations in the WSCT aligns with previous research that linked a oscillations to switching between brain states favoring visuospatial working memory stability or encoding flexibility (Ericson et al., 2025) and behavioral task switching in sensor-level EEG (Foxe et al., 2014; Verstraeten & Cluydts, 2002). Here, we demonstrate that local and large-scale α−β oscillations underlie the flexible switching between unstable cognitive representations of task rules. We speculate that this association was linked to individual reward sensitivity due to the strong association of α oscillations with engagement vs. disengagement found for visuospatial attention (Bonnefond & Jensen, 2025) putatively through their modulation of excitability (Iemi et al., 2017; Lange et al., 2013). Reward-related modulation of α oscillations was found in the ventromedial PFC that has earlier been shown to play a major role in reward processing and decision making (Domenech & Koechlin, 2015; Kennerley & Walton, 2011). Therefore, α−β band suppression over the PFC and PPC appear to improve reward-related cognitive flexibility through enhanced behavioral control.

In addition to local amplitude modulations, also fronto-parietal α desynchronization mediated by higher individual reward sensitivity predicted higher individual behavioral cognitive flexibility. This is in contrast with the association of large-scale increase in α−β synchronization with top-down attention (Doesburg et al., 2009; Lobier et al., 2018) and positive correlation with task performance (Palva et al., 2010; Sattelberger et al., 2024). We propose that in the context of the present task, disengagement of the α network serves as a reset of the internal top-down goal, i.e., the current cognitive rule representation. This is supported by some earlier studies in which α desynchronization during visual tasks was related to the tuning of attention in presence of competing sensory stimuli (Gross et al., 2004; Mylonas et al., 2016).

As a distinct neural sub-process of cognitive flexibility, we found a sustained increase in γ amplitudes that was associated with the processing of negative feedback. In line with γ amplitude being associated with punishment sensitivity, this modulation was found over the left cingulate cortex, which plays a central role in error monitoring (Carter et al., 1998; Walton et al., 2007). This suggests that γ amplitudes track the processing of errors and negative feedback in order to enable fast switching of cognitive representations based on contextual demands.

In contrast with a large body of EEG research showing that prefrontal and midline θ-band amplitudes enable cognitive flexibility control (Cooper et al., 2015; Duprez et al., 2020; Khan et al., 2025; Muralidharan et al., 2023; Riddle et al., 2020; Sauseng & Liesefeld, 2020), here, we found no evidence for θ amplitudes in enhancing behavior. In contrast, larger θ amplitudes predicted increased rule certainty across a trial series in our study and were explained by a tendency to anticipate rule changes. Our results thus indicate that increased θ amplitudes are not a facilitating mechanism for behavioral switching *per se* but rather characterize meta-contextual learning and anticipation of a new rule. Similarly, stronger prefrontal θ-band network synchronization was not directly linked to performance differences, but to a decreased reward and anticipation tendency, and to increased exploration tendency. Our results therefore provide new evidence that large-scale networks of prefrontal θ-band network synchronization might be a crucial neural inter-areal mechanism for rule exploration and internally guided flexible task-learning.

Particularly, the θ−γ PAC between the right dorsolateral PFC and bilateral parietal and visual cortices predicted better behavioral performance with fewer NLE, and shorter RT_shift_ and RT_stay_, While there is emerging evidence pointing out the significance of nested oscillations and specifically θ−γ coupling for cognitive functioning in attention and working memory (Brooks et al., 2020; Lisman & Jensen, 2013; Spooner et al., 2020) and global cognitive control (Riddle et al., 2021; Turi et al., 2020), we prove here for the first time its role in task switching. Such θ−γ PAC could allow for rapid communication between brain areas to implement pro- and reactive cognitive control (Pagnotta et al., 2024; Verguts, 2017; Voytek et al., 2015) and coordinate the top-down selection of the correct rule set in the WCST. This interpretation is supported by our finding that θ−γ PAC decreased over the course of a rule series.

## Conclusions

Our results demonstrate that the inter-individual variability in cognitive flexibility depends on differential sensitivity to feedback, which maps on to distinct oscillatory sub-processes. Our results therefore provide novel neural signatures for flexible switching between stable and unstable cognitive representations of contextual rules that predict the individual speed of rule learning.

## Methods and Materials

### Participants

Twenty-six healthy adults (mean age 33.2 (SD 10.2), age range 18–57 years, 11 females) participated in the study. All participants reported normal or corrected-to-normal vision, no recent history of neuropsychological or neurological illness, and physical eligibility to undergo MEG and MRI measurements. The study was approved by the Helsinki University Hospital ethical committee and was carried out according to the Declaration of Helsinki. All participants provided written informed consent before the experiment.

### Experimental task

Participants completed the Wisconsin card sorting test (WCST, Fig. 1A) by Berg (1948), adapted as a computerized version by Barcelo (2003) with the Presentation™ software (Neurobehavioral Systems, Inc., Albany, CA, USA). In the task, the participants needed to match cards according to three rule-categories: shape (circle, star, cross, or triangle), color (blue, yellow, red, or green), or number of items (one to four). The category by which to sort changed on every fifth trial and needed to be deduced by the participant through trial and error based on feedback. Participants were not informed how often the rule would change or what the next correct sorting category would be. In each trial, a sample card (bottom row), a black fixation cross, and four target cards (top row) were shown for 2.5 s before a white background. This was followed by a 1.5 s delay, during which the participant could match the stimulus card to one of the target cards, depending on the currently active rule (same number, color, or shape). A green or red feedback cross was shown for 0.25 s after the response to indicate a correct or incorrect response, respectively, followed by an inter-trial delay lasting 1.9 to 2.3 s. This delay served as a time window for rule selection, so that depending on the previous feedback, the participant would shift their internal rule representation or stay at the previously chosen rule. We used this rule-selection time window for our subsequent neural analysis.

The stimulus card was a rectangle of 7.3 x 5.5 deg, and the target cards were rectangles of 4.2 x 3.0 deg visual angle. A black fixation cross (0.65 x 0.48 deg) and black outlines of the stimulus and target cards were present throughout the whole trial in the center of this stimulus window. The target cards remained the same for the entire experiment. The sample card was selected randomly from 24 unambiguous cards of the original 64 WCST cards (Periánez et al., 2004), meaning that only one target card fit the current rule category. The rule orders were selected pseudo-randomly so that the same rule was not immediately repeated. The experiment consisted of three blocks with 150 trials each, where each block had an average duration of 15 minutes and participants could rest between blocks.

### Behavioral data analysis and the sequential learning model

After removal of trials without response (4 % of all trials), RTs were calculated as the duration from stimulus onset to response separately for trials following negative feedback (RT_shift_) and those following positive feedback (RT_stay_). Hit rates (HRs) were then calculated as the proportion of correct responses from all trials. We counted incorrect responses during the first two trials of a rule series as the number of early errors (NEE) and all subsequent errors as late errors (NLE), allowing us to assess the learning speed of participants across the trial set. To ensure that there was no bias in the response tendencies, we computed HRs for every trial in a rule series separately to confirm that the mean performance during early trials was at a chance level and exceeded thereafter. HRs, NLE, NEE were computed across all trials for each participant. To assess response speed, we computed mean RTs across trials for Shift (RT_shift_) and Stay (RT_stay_) trials.

For statistical comparisons of behavioral data, we used equivalent tests of both frequentist and Bayesian statistics (JASP Team, 2024). Statistical testing between the conditions (Shift vs. Stay) in RTs was performed with a two-tailed t-test for paired samples. To estimate the presence of a speed-accuracy trade-off, we computed the correlation of RT_shift_ and RT_stay_ with NLE across subjects using the Pearson correlation coefficient. To assess the co-variance of HR, NLE, RT_shift_, and RT_stay_ with age we computed the Pearson correlation coefficient with these measures and we likewise used independent-samples t-tests to estimate the influence of sex. For these tests, we reported the frequentist test parameters and also the Bayes Factors (BF10) as the likelihood of the independent variables to explain our data.

To further assess the individual factors underlying behavioral performance also at the single-trial level, we developed a sequential learning model to capture the effect of positive and negative feedback on task behavior (Fig. 1B). The model was based on the model by Bishara et al. (2010) and was used to estimate the subject parameters reward sensitivity ***r***, reflecting reward processing, and punishment sensitivity ***p***, measuring error monitoring, based on the single-trial behavioral data as in Bishara et al. (2010). In addition, as new behavioral measures, we estimated the subject parameters exploration tendency ***e*** and anticipation tendency ***a*** which assessed the effect of rule shifting despite preceding positive feedback and the subjects’ ability to predict an upcoming rule change after 5 trials, respectively. As our analysis revealed two outlier subjects with a punishment sensitivity ***p*** below 0.1 due to minimal error trials (Fig. 1I), we removed these subjects from further analysis of the model’s subject parameter data.

We then used multiple linear regression models to investigate if these parameters could explain differences in NLE, HRs and RTs and reported the Bayes Factor of the model (BFM) compared to the null model as well as each parameter’s Bayes Factor for model inclusion (BFi). Lastly, we also estimated the trial-to-trial rule certainty by creating an attention weight vector that represents the participant’s chosen rule in the previous trial (Bishara et al., 2010). Each value in this vector represents the probability of the active rule being shape, number, or color (e.g., [0,0,1] if the rule is known to be color). This vector is updated on every trial based on the card choice of the participant and the resulting feedback. Consequently, the rule uncertainty of one trial is first calculated as the Jensen-Shannon distance between the attention weight vectors of current trial and one trial before, being largest after a rule shift and becoming smaller after receiving positive feedbacks, from which we then obtain the rule certainty by inversion (rule certainty = 1 - rule uncertainty). The statistical significance of increased rule certainty across trials of a rule series was assessed with frequentist and Bayesian ANOVAs.

### MEG and MRI recordings, preprocessing and filtering

Participants sat approximately 57 cm from the screen (BenQ, height: 30 cm, width: 53 cm, resolution: 1920 × 1080, 60 Hz refresh rate). Stimuli were presented with the Presentation™ software (Neurobehavioral Systems, Inc., Albany, CA, USA) and a photodiode was used to achieve ± 1 ms precision of stimulus presentation. We recorded MEG with 204 planar gradiometers and 102 magnetometers with a Triux (Elekta Neuromag Ltd.) system at 1000 Hz sampling rate and an online band-pass filter of 0.03–330 Hz at the BioMag Laboratory in the Helsinki University Hospital. Ocular artifacts were measured with electro-oculogram (EOG) electrodes. Participants’ responses were recorded with a hand-held fiber-optic response device that responded to finger lifts.

In addition, whole head magnetic resonance images (MRIs; Siemens Verio 1.5 T) were taken from all subjects for the source-reconstruction of MEG data with the magnetization-prepared rapid acquisition gradient echo (MPRAGE) sequence at a resolution of 1 mm voxel size. Maxfilter software (Elekta Neuromag) was used for temporal signal space separation (tSSS) to suppress extra-cranial noise in MEG data, to adjust for head position changes during the measurements, and to interpolate bad channels (Taulu & Kajola, 2005). Head position changes over 10 mm were excluded from the raw data and raw MEG data was then notch-filtered at 50 Hz and harmonics. Independent component analysis (MATLAB) was used for identifying and extracting ocular and heartbeat artefacts. The MEG data was then frequency-filtered using 38 approximately logarithmically spaced Morlet wavelets from 3 to 143 Hz with m = 5 cycles.

### MEG source modelling

In order to obtain precise source localization of MEG signals optimized for every participant’s individual brain anatomy, we applied a multistep source modelling approach (Fig. 1C). The open-access software Freesurfer (http://surfer.nmr.mgh.harvard.edu) was used for volumetric segmentation of the MRI data, surface reconstruction, flattening, and cortical parcellation/labeling with the Destrieux atlas. The software MNE-Python (https://mne.tools/stable/index.html) was used for MEG-MRI colocalization, and preparation of the forward and inverse operators. For every subject, we created the source space with dipole orientations fixed to the pial surface normals and a 5 mm inter-dipole separation, a 1-layer boundary element method (BEM) model from the inner skull, and the BEM solution. To compute noise-covariance matrices (NCMs), we used a regularization constant of 0.05 and narrowband 151–299 Hz filtered resting-state data (1 min), which we recorded from each participant prior to the WCST in the same session. NCMs were then used to compute a forward solution and the inverse operator that transform the filtered complex single-trial MEG time series to source-vertex time series. Source-space vertices were collapsed into 400 cortical parcels using fidelity optimized collapse operators (Korhonen et al., 2014), assigning each parcel to one of 7 functional networks before morphing them into 200 larger parcels to increase statistical power.

### Data analysis

The time window of rule selection was used for all neural analysis. To extract this time window and an appropriately long baseline from the time series, we used 2.5 s epochs spanning from 0.55s prior to feedback onset to 1.95 s after. Single-subject oscillation amplitudes were computed for each wavelet frequency and for each parcel as an average across all trials of each condition (Shift and Stay), from which the baseline (0.5–0 s prior S1 onset) was then subtracted. Single-trial amplitudes were computed for the analysis of the trial order effect using the same approach but not averaging across the trials. To increase statistical power for condition-averaged and single-trial amplitude data, we averaged amplitudes within smaller time windows of 100 ms length that overlapped by 50 ms.

Phase-synchronization was computed for each wavelet frequency and between all parcels separately for both conditions and each subject. We first excluded 3 subjects with a small number of shift trials (N < 60) that did not yield sufficient effect sizes for the estimation of phase synchronization and equalized the number of trials between conditions to the lowest number within each subject as oscillation phase measures are sensitive to the number of trials. We then averaged the complex signal across trials for each condition and subject in smaller time windows of 400 ms length and 300 ms overlap, and computed the imaginary part of the phase locking value (iPLV) between each parcel-parcel pair for every time window. Single-trial phase synchronization was not estimated due to insufficient data points resulting from the file saving format. These data were baseline-corrected by subtracting the mean iPLV of the first time window (0.4–0 s prior S1 onset).

While using iPLV to estimate inter-areal synchronization excludes the direct effects of zero-phase lagged signal mixing, spurious interactions still remain (Palva et al., 2018). We therefore removed edges (pairwise connections) between parcels for which the source reconstruction accuracy, fidelity, was below 0.1 (3.5 % of parcels). To further exclude spurious connections, we removed edges between parcels that exhibited the greatest signal leakage with their neighbors, measured with cross-patch PLV (fidelity radius greater than 0.4). A total of 17.8 % of all possible edges were thus excluded from the analyses. This created adjacency matrices for each frequency and for each subject that defined a graph made up of nodes and edges, where nodes are cortical parcels and edges are the connections between nodes.

To then estimate n:m phase amplitude coupling (PAC), we compared the phase of all low frequencies (3– 8 Hz) to the phase of the amplitude envelope of high frequencies (9–143 Hz) by computing the PLV between all edge pairs. This was done for each subject and for the averaged trials of conditions Shift and Stay separately, in the time window 0.6 to 1.6 s after feedback onset. Because PLV is sensitive to the number of trials, we equalized the number of trials between conditions to the lowest number within each subject before averaging, again excluding the 3 subjects with N < 60 Shift trials.

### Statistical analysis of neuronal data

Group-level statistical analyses of neuronal effects were then carried out. For oscillation amplitudes, statistical analysis were carried out separately for each parcel, time window and frequency across subjects. For inter-areal phase-synchronization and PAC, the statistical analysis was carried out per each edge (parcel-parcel connection), time window (only one for PAC) and frequency across subjects.

A two-sided Wilcoxon-signed rank test at α = 0.05 was used for testing statistical differences between either condition (Shift and Stay) and the baseline, as well as their contrast, Shift-Stay. To estimate statistically significant correlations of oscillation amplitudes, inter-areal synchronization, and PAC with NLE, RT_shift_, and RT_stay_, we calculated Spearman’s r at α = 0.05 for every frequency, time window and parcel or edge. To estimate the effect of trial series on single-trial amplitudes, Spearman’s r at α = 0.05 was also used to compute statistically significant changes in oscillation amplitudes across a trial series for every frequency, time window and parcel. We furthermore calculated Spearman’s r at α = 0.05 to assess the correlation of the model subject parameters with the condition-averaged amplitude and phase synchronization across subjects, separately for every frequency, time window and parcel or edge.

To estimate the robustness of oscillation amplitude modulations and that of changes in inter-areal phase synchronization across the whole cortex, we calculated the fraction of cortical parcels with a significant effect out of all 200 parcels (*P*) and similarly, the connection density (*K*), which is the fraction of significant edges out of all 39,800 edges. For phase synchrony, statistical thresholding was further used to construct two group-level adjacency matrices that contained the average iPLV of significant edges across time and frequencies where the non-significant values were set to zero, for the θ- and α-band.

When analysing statistical significance of oscillation amplitude and phase synchrony effects, we then removed false positives caused by multiple statistical tests by pooling all significant observations over all parcels for amplitude analysis and edges for synchrony and then discarded a percentage of the least-significant comparisons predicted to be false discoveries equivalent to the α level (Palva et al., 2010; Siebenhühner et al., 2020). To remove any remaining false positives, a threshold q was defined for the number of significant observations that could arise by chance in any of the frequencies even after controlling for multiple comparisons (Puoliväli et al., 2020). This was estimated by simulating random processes at the null hypothesis, using the same number of tests performed in the actual experiment, and recording the residual fractions of significant observations after the elimination of the number of significant observations predicted to be false positives by the α level. Given the number of cortical parcels and time windows in which statistical analyses were performed for each condition, q was estimated to correspond to 8 % of parcels and 0.67 % edge density and was removed from the fraction of significant parcels (*P*) or the connection density (*K*). We only visualized oscillation amplitude modulations and inter-areal synchronization changes that exceeded the q threshold across frequencies in time-frequency representations (TFRs).

For statistical analysis of PAC at the group level, we estimated the statistical significance of each pairwise interaction for each of the 1:m ratios. We identified interactions that were significantly stronger or weaker in the Shift and Stay conditions than at baseline by computing time-shifted surrogate data and deeming connections with a ratio of 2.42 (corresponding to p < 0.01) or higher as significant. Spurious connections were removed by estimating amplitude coupling between low and high frequencies and identifying triangle motifs (Siebenhühner et al., 2020), after which we removed edges below the q threshold of 3.5 % as described above. Shift- and Stay-condition PAC were contrasted using a two-sided Wilcoxon signed-rank test with a statistical significance level p < 0.05 and corrected for multiple comparisons by removing the 5 % largest p-values as well as edges below the q threshold of 0.39 %. The trial order effect and behavioral correlations were again corrected for any remaining false positives by applying the thresholding with q = 0.28 %.

Post-hoc analyses were carried out on time-frequency (TF) windows of interest that were selected to contain the most robust (largest P values for amplitude and K for phase synchronization) significant neuronal effects. This includes the estimation of effect sizes, for which we calculated the average difference between all parcels/edges within the described TF windows of interest as r = Z √ N (Rosenthal, 1991). We also obtained the mean graph strength (mean GS) of the averaged edges within functional networks in the TF windows of interest of the θ- and α-band from the thresholded adjacency matrix (Fig. 3D).

### Graph visualization

The spectro-temporal patterns of phase synchrony and PAC modulations were first visualized as TFRs, in which we identified the frequency bands and TF windows with the most robust effects. We then showed the most important hubs and connections in those networks as graphs by visualizing the edges that were significant in more than 15% of the TF bins across the selected time and frequencies of the thresholded adjacency matrix.

Node centrality metrics were further used to identify highly connected nodes in the graphs that putatively play a key role in network communication. Here, we used the graph metric *Degree,* that is the number of edges of a node to other nodes, to select the most connected nodes which act as communication hubs in the network. We then visualized graphs that contained the nodes that were within the 65th percentile of largest-degrees and their 200 strongest edges, i.e., the edges with the highest iPLV for phase synchrony and PLV for PAC. To further mitigate the contribution of spurious interactions caused by the concurrent presence of true interactions and linear mixing, a hyperedge-bundling approach was used in the visualization (Wang et al., 2018). The hyper-edges thus bundled together edges that putatively originated from a single true edge among the spurious edges caused by source-leakage (Palva et al., 2018).

## CRediT author contribution(s)

JuS: Conceptualization, Data curation, Formal analysis, Investigation, Methodology, Project administration, Software, Visualization, Writing –original draft, HH: Conceptualization, Methodology, ML: Formal analysis, Methodology, JaS: Data curation, Investigation, Software, JMP: Conceptualization, Funding acquisition, Methodology, Resources, Writing – review & editing, MMH: Conceptualization, Methodology, Writing – Review & Editing, SP: Conceptualization, Funding acquisition, Methodology, Project administration, Supervision, Visualization, Writing – original draft, Writing – review & editing

## Acknowledgements

The authors would like to thank Jasmin Elonen for preprocessing the MEG data.

