## Supplementary Figures for "Distinct neural oscillations predict covert switching between stable and unstable cognitive representations"

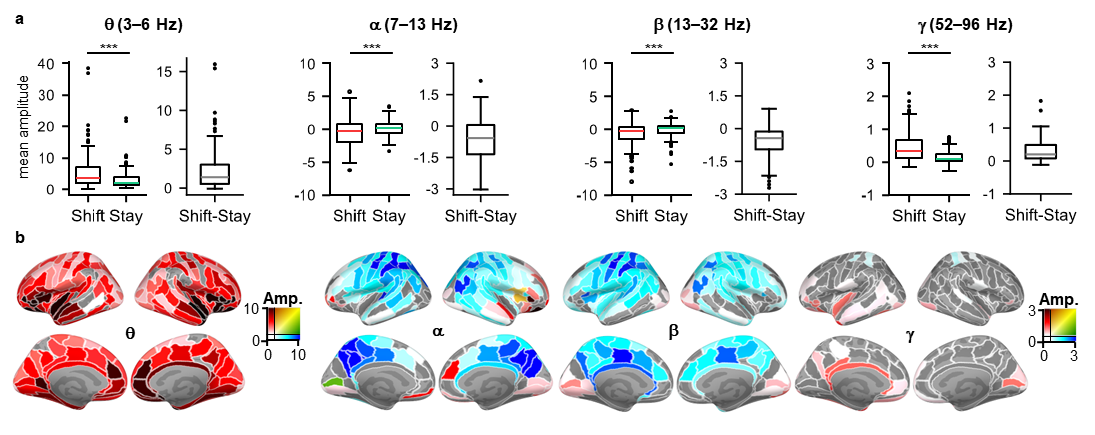
**Supplementary Figure 1. Effect sizes and cortical localization of Shift-Stay amplitude differences. a.** Boxplots indicating the mean parcel amplitudes in Shift and Stay trials for all frequency bands with a significant effect (θ−β: 0.8-1.5 s; γ: 0.9-1.2 s) and their differences. Asterisks indicate significant differences between conditions. **b.** Inflated brain plots showing the mean amplitude of parcels with significant oscillation amplitude changes between Shift and Stay trials in the same TF windows of interest as in **a**.

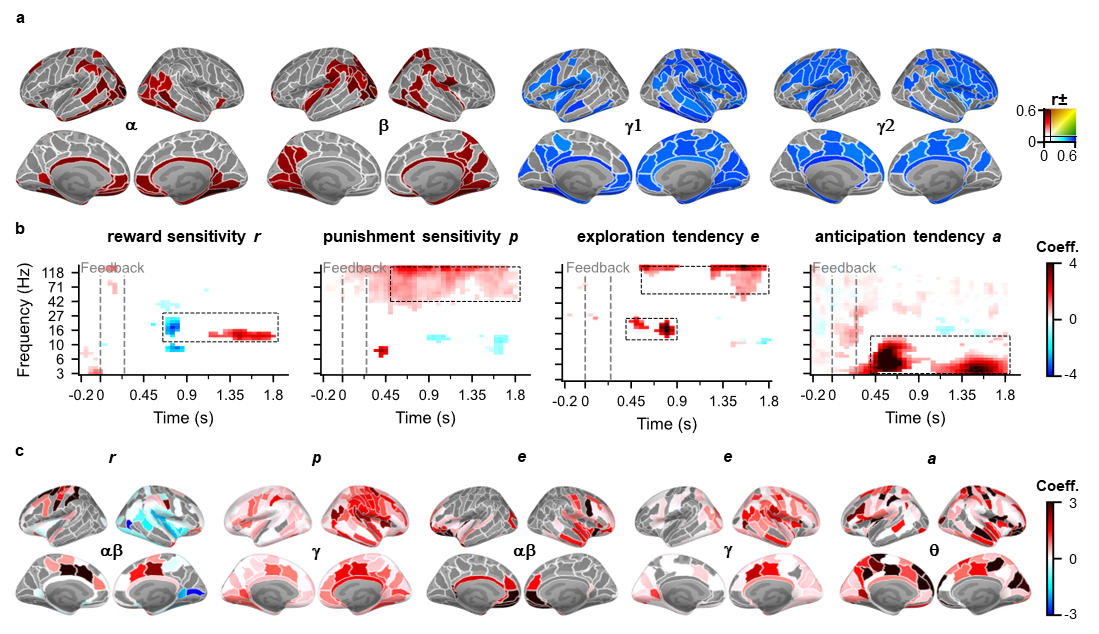
**Supplementary Figure 2. Amplitude correlations with performance and effects of the subject parameters. a.** Brain anatomies showing the parcels with a significant correlation with number of late errors (α−γ1) and RT_stay_ (γ2). Color indicates the strength of the correlation coefficient (Coeff.). **b.** Model fit for each subject parameter with the mean amplitudes of Shift and Stay trials. The color shade expresses the coefficients of parcels with a significant model fit (Linear Regression, p < 0.05, FDR corrected, q = 0.08; N = 24). Dotted boxes indicate the TF windows of interest for a frequency band. **c.** Inflated brain plots showing the coefficients of parcels with significant model fit by subject parameters in the TF windows of interest marked in **b**.

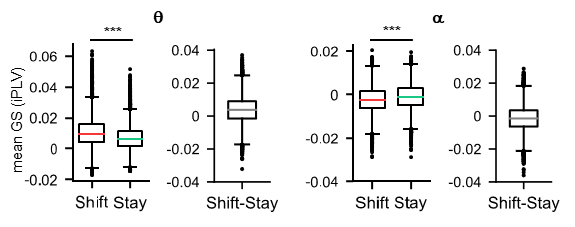
**Supplementary Figure 3. Effect sizes of phase synchrony (iPLV) results for Shift and Stay conditions.** Boxplots showing the effect sizes in the θ- and α-bands of the difference between Shift and Stay trials, the asterisk indicating a highly significant difference (Wilcoxon signed-rank test, *p < 0.05, **p < 0.01, ***p < 0.001).

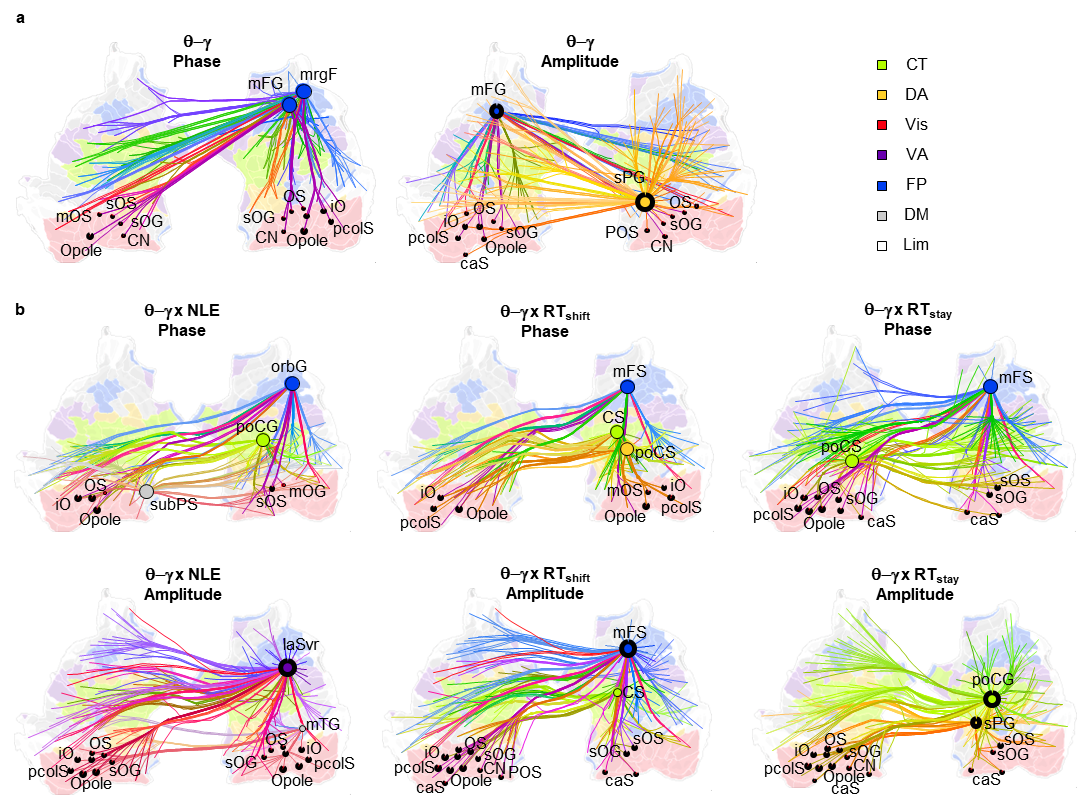
**Supplementary Figure 4**. θ−γ PAC network underlying cognitive flexibility. **a.** Graphs showing the significant θ−γ PAC networks by largest degree of phase to amplitude coupling (left) and amplitude to phase coupling (right). **b.** Graphs of θ−γ PAC by behavioral measure, showing the significant edges with the largest degree of coupling of phase to amplitude (top) and of amplitude to phase (bottom).
